# NanoNet: Rapid end-to-end nanobody modeling by deep learning at sub angstrom resolution

**DOI:** 10.1101/2021.08.03.454917

**Authors:** Tomer Cohen, Matan Halfon, Dina Schneidman-Duhovny

## Abstract

Antibodies are a rapidly growing class of therapeutics. Recently, single domain camelid VHH antibodies, and their recognition nanobody domain (Nb) appeared as a cost-effective highly stable alternative to full-length antibodies. There is a growing need for high-throughput epitope mapping based on accurate structural modeling of the variable domains that share a common fold and differ in the Complementarity Determining Regions (CDRs). We develop a deep learning end-to-end model, NanoNet, that given a sequence directly produces the 3D coordinates of the Cɑ atoms of the entire VH domain. For the Nb test set, NanoNet achieves 1.7Å overall average RMSD and 3.0Å average RMSD for the most variable CDR3 loops. The accuracy for antibody VH domains is even higher: overall average RMSD < 1Å and 2.2Å RMSD for CDR3. NanoNet runtimes allow generation of ~1M nanobody structures in less than an hour on a standard CPU computer enabling high-throughput structure modeling.

## Introduction

The large and diverse repertoire of the immune receptors, including antibodies and T cell receptors (TCRs) is behind the specific antigen recognition mechanism ^1^. Next generation sequencing provides a glimpse into the blood circulating repertoires. However, the antigens and the epitopes remain unidentified. Moreover, antibodies are the most rapidly growing class of human therapeutics for a range of diseases, including cancer or viral infections. Despite their successful application, there are challenges in high-throughput cost-effective manufacturing of monoclonal antibodies (mAbs), as well as intravenous administration route. Nanobodies (Nbs) are small and highly-stable fragments derived from camelid heavy chain only antibodies ^2,3^. They can reach binding affinities comparable to antibodies. Nbs can be manufactured easily in microbes and administered by aerosolization ^4^. Rapid Nb development is possible by camelid immunization ^5,6^ or synthetic design and screening ^7^.

Epitope characterization is an important part of therapeutic antibody (mAb or Nb) discovery. It is critical to select epitope specific sequences from a large pool of candidates. However, high-throughput experimental structural characterization of hundreds or thousands of antibody-antigen complexes remains challenging. Computational methods for modeling antibody-antigen structures from individual components frequently suffer from high false positive rate, rarely resulting in a unique solution. There are two main bottlenecks: low accuracy of antibody CDR loop modeling and antibody-antigen scoring functions.

Antibody modeling most often proceeds in two steps. First, the conserved framework region is modeled by comparative modeling. Second, the variable CDR loops are modeled. The CDR3 loop which is highly variable and long presents a mini *ab initio* folding problem. While there are existing tools for mAb and TCR modeling ^8–10^, dedicated algorithms for reliable Nb modeling are unavailable. Compared to mAbs, Nbs generally have longer CDR3 loops and are devoid of light chains, adding additional degrees of freedom for accurate loop modeling.

Recently, deep learning has been successful in addressing challenging and fundamentally important questions in structural biology, including protein folding and *de novo* protein design ^11–15^. Moreover, deep learning was successful in predicting restraints for the mAb CDR3 heavy chain loop modeling ^16,17^. All the deep learning-based algorithms, with the exception of novel AlphaFold2 and RosettaFold ^14,15^, use deep learning for restraints generation, requiring an additional optimization step to generate 3D structures. The structure generation step is time consuming. For example, RosettaAntibody requires about 30 minutes per model, where ~50 models are generated per single sequence.

Here we use deep learning for accurate end-to-end prediction of Nb structures. While our main goal is accurate Nb modeling, NanoNet can also accurately model VH domains of the antibodies and Vβ domains of TCRs. Our deep learning model accepts the sequence (Nb, mAb VH domain, or TCR Vβ domain) as an input and produces coordinates of the Cɑ atoms. NanoNet improves upon existing models using direct end-to-end learning that enables the network to learn the full 3D structure without dividing the modeling problem into framework and CDRs modeling.

## Results

### Summary of the method

The input to the NanoNet is the sequence (mAb VH, Nb, or TCR Vβ domains) and the output is the Cɑ coordinates for the input sequence. The network was trained on a dataset of ~2,000 heavy chains of mAbs and Nb structures. The framework region of the antibodies is highly conserved with Cɑ RMSD under 1Å between aligned structures. Therefore, we achieved transformational invariance for predicting 3D coordinates by aligning all the structures of the training set on a randomly selected reference structure. The VH domain structures were aligned using MultiProt algorithm with order-dependence and accuracy of 0.8Å ^18^ (Fig. 1B). This structure alignment enables the network to directly learn the VH domain 3D structure. The network is a convolutional neural network (CNN) that consists of 2 blocks of 1D Residual Neural Networks (ResNet) ^19^ (Fig. 1A). The loss is defined as an MSE (Mean Squared Error) on the Cɑ coordinates, which is equivalent to the squared RMSD, and an additional term that optimizes the distance between consecutive Cɑ atoms to 3.8Å.

**Figure 1:**
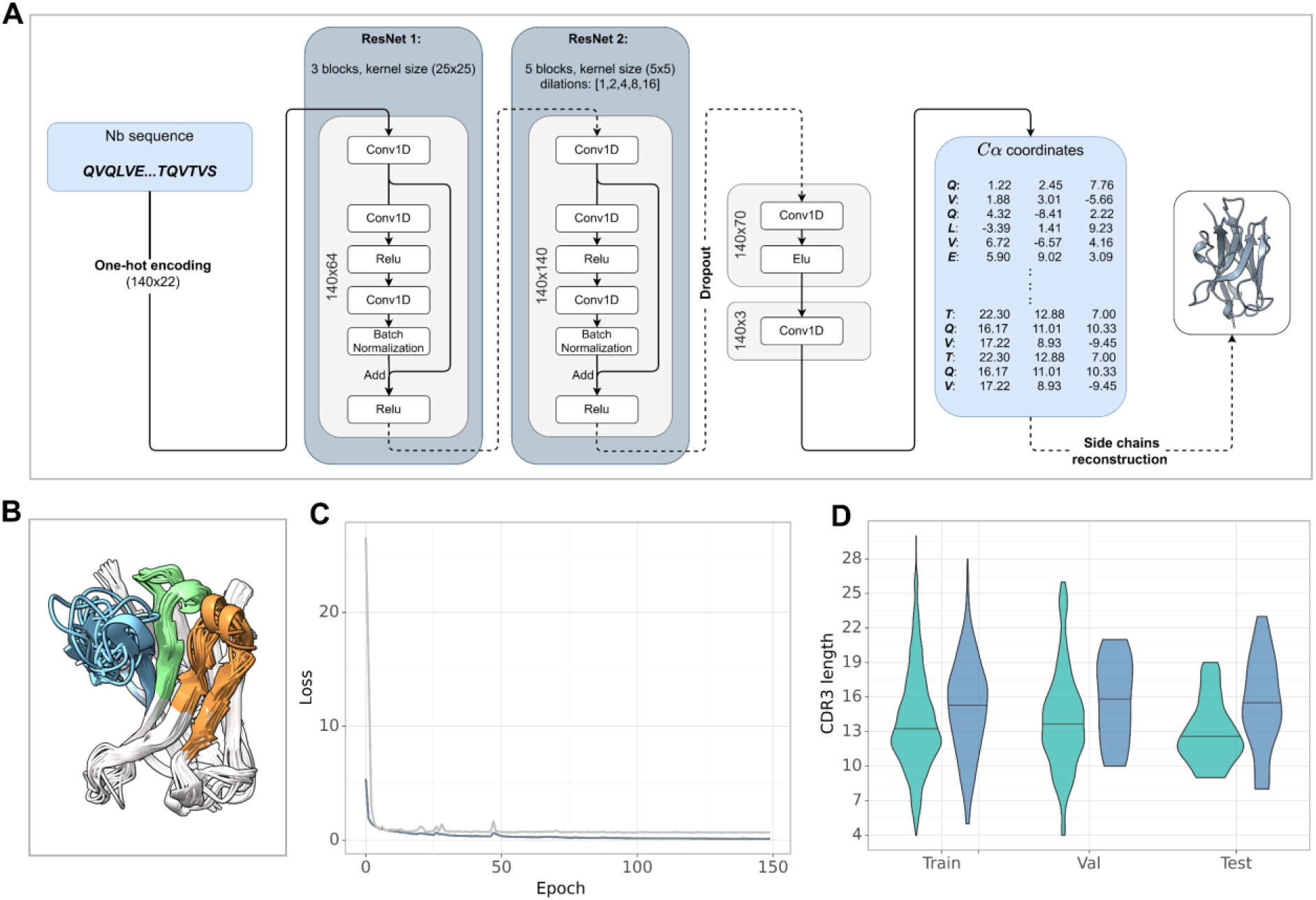
A. NanoNet architecture: the input of the network is the one-hot encoding of the sequence that goes into two 1D ResNets. The output is the Cɑ coordinates. B. Structural alignment of the antibodies from the training set prior to coordinate extraction for training the network. C. Training and validation loss during the training process. D. CDR3 lengths of the training, validation, and test sets (mAb green, Nb blue).

### NanoNet produces high-accuracy Nb models

The overall average RMSD on the Nb_test (Methods) is 1.7Å and the median is 1.5Å (Fig. 2A, Fig. S2A, Table S1). The CDR1 and CDR2 loops are also accurately modeled with average RMSDs of 2.4Å and 1.7Å (median 2.3Å and 1.5Å), respectively. For the most challenging CDR3 loop, we obtain an average RMSD of 3.0Å and a median of 2.7Å. NanoNet obtains highly accurate models also for Nbs with longer CDR3 loops (> 12 amino acids) (Fig. S2B). Due to their longer length compared to mAbs (Fig. 1D), Nb CDR3 loops often contain short 3_10_ helices. We find that NanoNet accurately reproduces such secondary structures in long CDR3 loops (Fig. 2C, PDB 6xxo, loop length 17 residues). We also manually examine cases where NanoNet produces higher RMSD and find that often the loop conformation is correct but it’s orientation with respect to the frame is shifted (Fig. 2C, PDB 7n0r).

**Figure 2:**
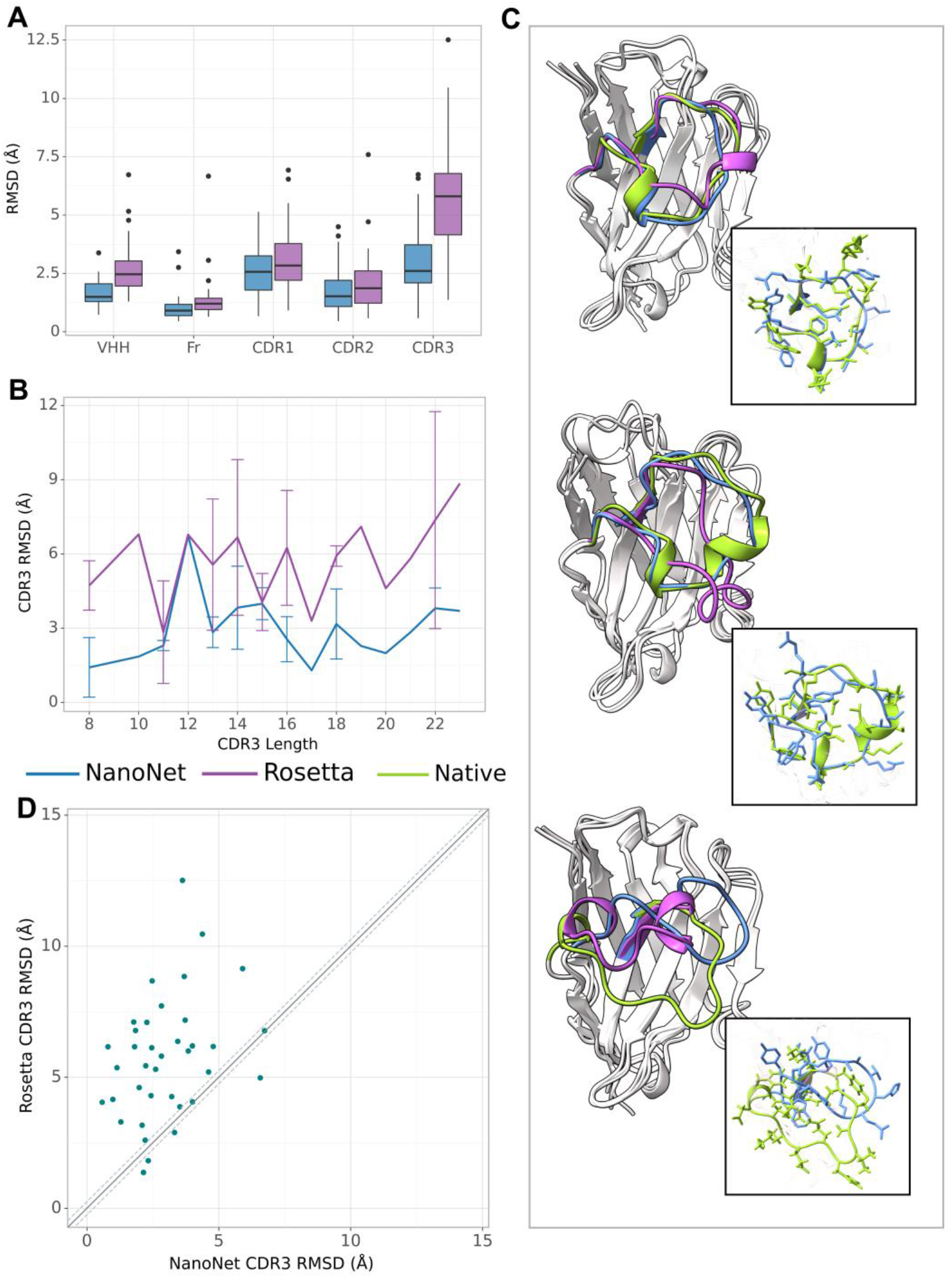
Nb modeling results. A. Boxplots of RMSDs of the whole VHH region, framework, CDR1-3 loops for the test set Nb. B. RMSD of CDR3 loop as a function of loop length on the test set of 37 Nbs. C. Test set examples of modeled structures by NanoNet (blue), RosettaAntibody (purple) vs. experimental (green): PDB 6xxo (top), 6lr7 (middle) and 7n0r (bottom), RMSDs 1.60Å, 1.07Å and 2.53Å, for NanoNet, and 2.16Å, 2.21Å and 1.94Å for RosettaAntibody, respectively. D. CDR3 loop RMSD for NanoNet vs. RosettaAntibody, each dot represents a structure from the test set. The dotted line corresponds to 0.25Å RMSD.

We compare our results to the results of RosettaAntibody (Fig. 2A-D). RosettaAntibody failed to generate models for 7 Nbs from the Nb_test due to the lack of suitable templates (required for CDR1-2 modeling) or program failures. For additional 6 Nbs manual CDRs definition was necessary to produce models. Overall, NanoNet has lower RMSD for 29 out of 37 test cases and equal RMSD for 4 test cases (Fig. 2D). The CDR3 RMSD is almost twice more accurate (3.0Å vs. 5.7Å, 47% improvement), while there is also improvement in the overall RMSD (1.7Å vs. 2.7Å).

### NanoNet VH models are comparable to state-of-the-art mAb modeling approaches

We test the method on VH modeling using the mAB_test (Methods). Overall, due to shorter average loop length and a larger number of mAbs in the training set, we obtain highly accurate models with average sub angstrom accuracy of 0.99Å overall RMSD (median 0.92Å). The CDR1, CDR2, CDR3 loops are also highly accurate with mean RMSDs of 0.82Å, 0.83Å, 2.25Å (medians 0.55Å, 0.60Å and 2.04Å), respectively (Fig. 3A-B, Table S2). Similarly to Nb modeling, NanoNet models have significantly higher accuracy compared to RosettaAntibody (Fig. 3A-D). NanoNet results are comparable to the accuracy reported for DeepH3 and DeepAb ^16,17^ (Fig 3A-C) with slightly lower number of outliers with high CDR3 RMSD for NanoNet (Fig. 3E). We find that NanoNet can reproduce short secondary structure motifs in the CDR3 loops. For example, we can reconstruct the β turn in the CDR3 loop consisting of 14 residues (PDB 1jfq, Fig. 3C). The main advantage of NanoNet compared to other deep learning models, such as DeepH3 or DeepAb, is that it directly produces the structure in millisecond time frame and therefore is applicable to high-throughput modeling of large databases. For comparison, loop optimization with RosettaAntibody takes at least 30 minutes per loop (with or without the distance and angle restraints from deep learning) and at least 50 loop conformations are usually generated per antibody.

**Figure 3:**
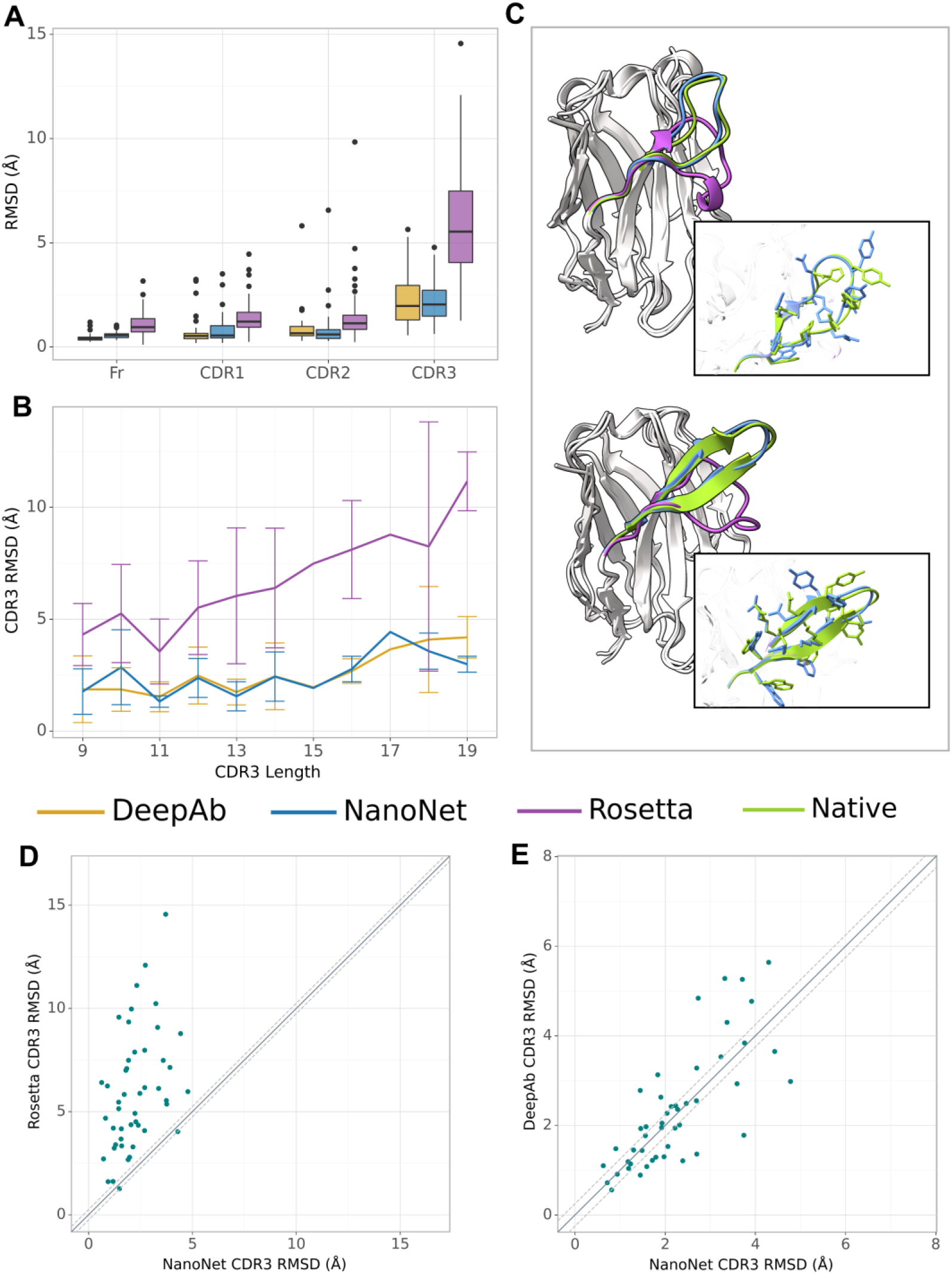
mAb VH modeling. A. Boxplots of RMSDs of the framework, CDR1-3 loops for the test set mAbs. B. RMSD of CDR3 loop as a function of loop length on the test set of 47 mAbs. C. Test set examples of modeled structures by NanoNet (blue), RosettaAntibody (purple) vs. experimental (green): PDB 3t65 (top) and 1jfq (bottom), RMSD 0.46Å and 0.73Å, for NanoNet, and 2.30Å and 2.57Å for RosettaAntibody, respectively. D. CDR3 loop RMSD for NanoNet vs. RosettaAntibody, each dot represents a structure from the test set. The dotted line corresponds to 0.25Å RMSD. E. same as D for NanoNet vs. DeepAb.

### Effect of sequence identity cut-off used for train and test set separation

Due to high baseline sequence identity between the antibodies (minimum 75% sequence identity, average 88% for mAbs and 84% for Nbs), splitting the input structures into train and test sets needs to be done carefully to prevent overfitting. To obtain the most optimal split cut-off, we compare the performance of NanoNet for each test set case as a function of maximal sequence identity (MSI) in the training set. We find that for the mAb_test there is a significant negative correlation of RMSD with the training set MSI (Fig. 4A-B). This can be explained by the high split cut-off of 99% for the RosettaAntibody dataset that was also used for training of DeepH3 and DeepAb models. In contrast, for the Nb_test that was generated using the 90% cutt-off, there is no correlation between RMSD and MSI (Fig. 4C-D). Similar results were obtained for TCRs (Fig. S1C-D).

**Fig 4:**
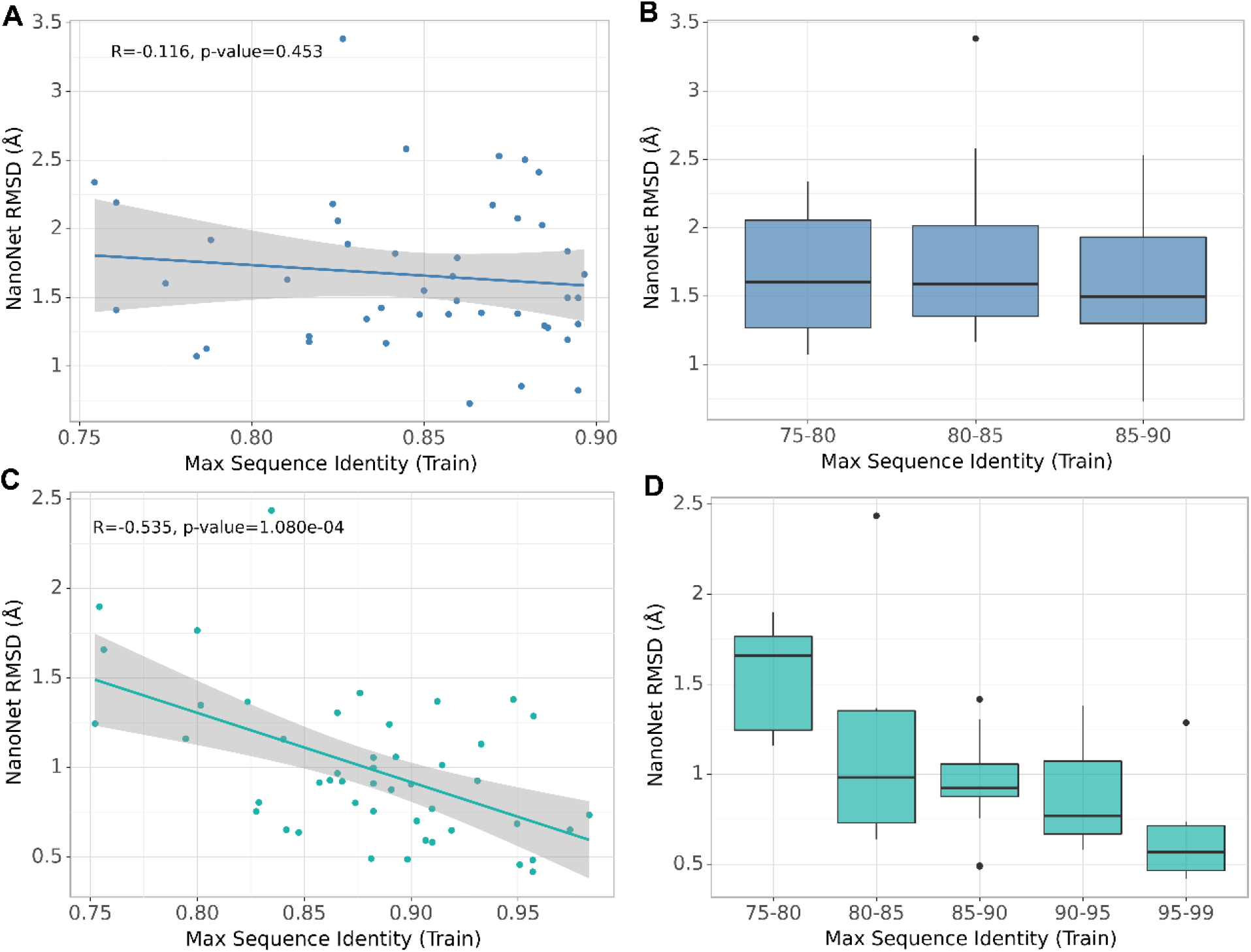
Effect of splitting the train and test sets using different sequence identity cutoffs. A and C. Test set performance (measured as VH RMSD) as a function of maximal sequence identity to the train set, each dot represents a structure from the test set for NanoNet data split (A) and DeepAb/RosettaAntibody split (C). B and D. Test set performance with boxplots for sequence identity ranges for NanoNet split (B) and DeepAb split (D).

### Retraining for TCR Vβ modeling

The PDB contains only 200 non-redundant TCR structures (using 99% sequence identity cut-off), a number that is too low for training a deep network. Because the TCRs are structurally similar to the antibodies (Fig. S3A), we tested if transfer learning is applicable by re-training the NanoNet on the TCR structures dataset. Our test set contains 15 structures of variable CDR3 length (Fig. S3B). Despite the small number of available structures, accurate TCR models can be predicted for the frame and CDR loops. We obtained a mean and median RMSD values of 1.30Å, 1.05Å for the entire Vβ and mean RMSD values of 0.77Å, 1.19Å, 2.05Å for CDR1, CDR2, CDR3 (medians 0.65Å, 0.80Å and 2.02Å), respectively (Fig. S3C). We suggest that the accuracy will improve upon availability of additional TCR structures.

### Antigen-Nb docking with NanoNet models

To further test the performance of NanoNet in the context of epitope prediction, we docked the modeled nanobodies from the test set to the antigens with known antigen-Nb structures. Antigen-Nb docking is highly challenging because docking algorithms have difficulties in sampling docked configurations with inaccurate CDR loops conformations. Additional difficulty is ranking the correct models among top-scoring ones. Here we tested whether the accuracy of NanoNet Nb models is sufficient to enable sampling of the correct orientation. Docking was applied for 17 Nbs (7 Nbs that interact with SARS-CoV-2 spike RBD ^20–24^, 2 Nbs that bind SARS-CoV-2 N protein ^25^, 1 Nb binding to SARS-CoV-1 spike RBD, 1 Nb binding to the Ebola RNA methyltransferase ^26^, and 4 MNV capsid protein P-domain Nbs ^27^). In addition, we docked two high affinity SARS-CoV-2 RBD Nbs (Nb21, Nb105) that bind to different epitopes^28^. The docking algorithm was able to generate an acceptable quality model according to the CAPRI evaluation criteria (ligand RMSD < 10Å or interface RMSD < 4Å) for all the structures predicted by NanoNet. Overall, we achieved a mean minimal ligand RMSD of 5.50Å and a mean minimal interface RMSD of 3.43Å (Table S4). Medium quality models (ligand RMSD < 5 Å or interface RMSD < 2) were sampled for 8 out of the 17 Nb structures. These results suggests that NanoNet generated models can be used in docking for epitope mapping.

### Correlating sequence and structure similarity using NanoNet

Next, we use a large dataset of high affinity Glutathione S-transferase (GST) binding Nbs ^29^ to explore the sequence-structure similarity relationship. We hypothesize that Nbs with similar sequences will have similar structures. The dataset contains 6,222 sequences with 1,476 distinct CDR3s and 2,566 distinct CDR1-3 combinations. Because the Nbs were obtained from a single llama using a novel proteomic platform ^29^, it contains diverse clusters of Nbs with high sequence similarity within the cluster. We used NanoNet to model all the 6,222 Nbs in a matter of seconds. We then quantify the structural similarity for all pairs of Nbs and produce a distance matrix. The structural distance between a pair of Nbs was defined using the number of corresponding Cɑ atoms pairs (one from each structure) with distance < 1Å (Methods). We used the pairwise distance matrix to cluster the structures by K-Medoids algorithm ^30^ with k=12. To compare these structural clusters with the sequence similarity we apply 2D dimensionality reduction of the sequences using t-SNE ^31^ on the multiple sequence alignment of the Nbs (Methods). To analyze sequence-structure similarity between the Nbs, we colored each sequence in the t-SNE sequence embedding by the structural cluster number (Fig. 5A) as well as the CDR3 length (Fig. 5B). We find that the structural clusters map well onto the sequence clusters. The clusters contain tens to hundreds Nb sequences and most likely represent the affinity maturation process where residue substitutions are made to improve stability and antigen binding affinity while the Nb structure is maintained (Fig. 5C-D). We also find several outlier Nbs that are far from their structural cluster in the sequence space, indicating that the specific mutations may change the Nb conformation. The correlation between structure and sequence clustering is independent of CDR3 length (Fig. 5B).

**Fig. 5:**
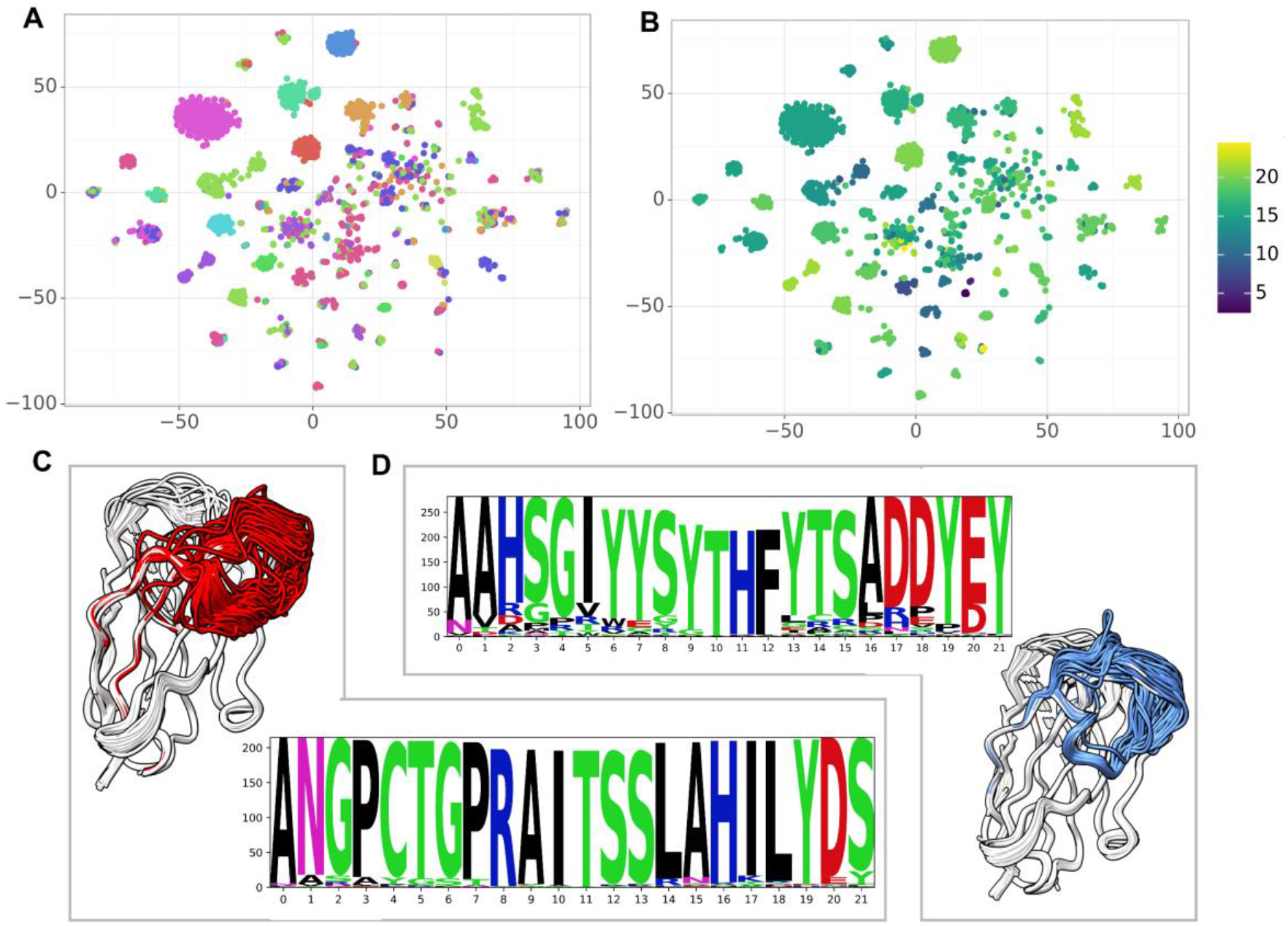
Correlating sequence and structure similarity. A. The t-SNE sequence embedding colored by the structural cluster number (12 clusters). B. The t-SNE sequence embedding colored by the CDR3 length. C-D. Examples of structural clusters and their MSA, CDR3 is colored by the cluster color in the t-SNE sequence embedding.

### Comparison to AlphaFold2

Publication of the highly accurate structure prediction model ^14^ prompted us to explore the accuracy of Nb structures as predicted by AlphaFold2. Direct comparison to AlphaFold2 is not possible because we do not know which PDB structures were used in the training of the model. We generated models for several Nbs published in 2021 assuming that they were not part of the training set. In total, 16 Nbs from the test set were compared to AlphaFold2. The CDR3 RMSD was comparable for 4 cases, NanoNet had lower CDR3 RMSD in 7 cases, and AlphaFold2 had lower RMSD in 5 cases (Fig. 6A). NanoNet achieved mean VHH RMSD and mean CDR3 RMSD values of 1.57Å and 2.69Å respectively compared to 2.04Å and 3.23Å for AlphaFold2 (Fig. 6B, Table S3). In most cases NanoNet and AlphaFold2 produce relatively similar CDR3 conformations, but we find that in some cases AlphaFold2 generated models does not reproduce the Nb framework accurately (Fig. 6C). In cases where NanoNet achieved relatively high CDR3 RMSD, AlphaFold2 predictions were similar to NanoNet, however both were far from the crystal structure (Fig. 6D). We suggest this happens due to the conformational changes upon antigen binding. These examples highlight the advantage of Nb specific structure prediction model in speed and accuracy.

**Fig. 6:**
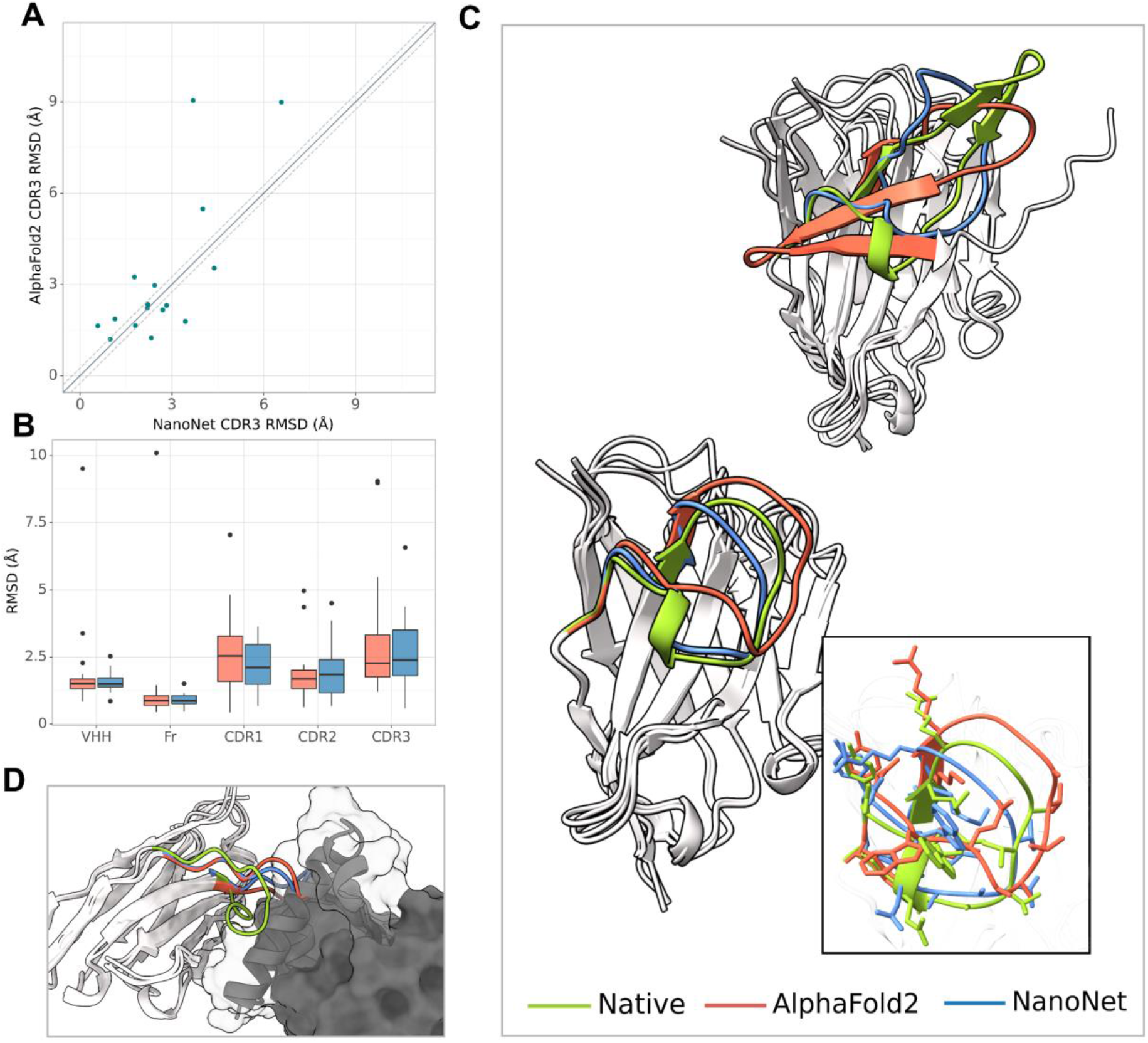
Comparison to AlphaFold2. A. CDR3 loop RMSD for NanoNet vs. AlphaFold2, each dot represents a structure from the 16 Nbs test set. B. Boxplots of RMSDs for the whole VHH region, framework, CDR1-3 loops for the 16 Nbs. C. Test set examples of modeled structures by NanoNet (blue), AlphaFold2 (red) vs. experimental (green): PDB 7ldj (top, CDR3 length - 23) and 7lvu (bottom, CDR3 length - 16), RMSDs 2.02Å, 1.37Å, for NanoNet and 9.51Å,1.58Å for AlphaFold2, respectively. D. Nb (PDB 6xzu) docked to its antigen, native structure –green, AlphaFold2 - red, NanoNet - blue.

**Table 1:** summary of mean RMSD results for the different test sets. Top - Nb test set, middle-mAb test set, bottom - 16 Nbs from the Nb test set (released in 2021).

| Method | VHH | Fr | CDR1 | CDR2 | CDR3 |
| --- | --- | --- | --- | --- | --- |
| **RosettaAntibody** | 2.71 ($\pm 1.13$) | 1.39 ($\pm 1.01$) | 3.09 ($\pm 1.37$) | 2.13 ($\pm 1.29$) | 5.73 ($\pm 2.33$) |
| **NanoNet** | 1.68 ($\pm 0.57$) | 1.02 ($\pm 0.57$) | 2.54 ($\pm 1.05$) | 1.70 ($\pm 0.98$) | 2.99 ($\pm 1.48$) |

| Method | Fr | CDR1 | CDR2 | CDR3 |
| --- | --- | --- | --- | --- |
| **RosettaAntibody** | 1.08 ($\pm 0.58$) | 1.45 ($\pm 0.84$) | 1.53 ($\pm 1.54$) | 5.95 ($\pm 2.88$) |
| **DeepAB** | 0.43 ($\pm 0.18$) | 0.72 ($\pm 0.66$) | 0.85 ($\pm 0.81$) | 2.33 ($\pm 1.32$) |
| **NanoNet** | 0.57 ($\pm 0.18$) | 0.82 ($\pm 0.66$) | 0.83 ($\pm 0.96$) | 2.25 ($\pm 1.05$) |

| Method | VHH | Fr | CDR1 | CDR2 | CDR3 |
| --- | --- | --- | --- | --- | --- |
| **AlphaFold2** | 2.04 ($\pm 2.09$) | 1.45 ($\pm 2.32$) | 2.63 ($\pm 1.68$) | 1.88 ($\pm 1.19$) | 3.23 ($\pm 2.49$) |
| **NanoNet** | 1.57 ($\pm 0.41$) | 0.91 ($\pm 0.24$) | 2.27 ($\pm 0.90$) | 1.96 ($\pm 1.08$) | 2.69 ($\pm 1.49$) |

## Discussion

We have developed a high-throughput accurate end-to-end deep learning-based method for 3D Nb modeling. Compared to previous antibody modeling approaches, the method directly predicts the 3D coordinates for the whole Nb sequence without separate modeling of frame and CDR regions. Because NanoNet was trained on antibody VH domain structures (as well as Nbs), it has subangstrom accuracy in VH modeling. The accuracy of the method is significantly higher than standard loop modeling methods and comparable to deep learning based approaches (Figs. 2,3), including AlphaFold2 (Fig. 6). We find that despite longer CDR3 loops of the Nbs, NanoNet can model them with high accuracy, reproducing the short secondary structures, such as 3-10 helix or beta-turn (Figs. 2,3). We extend the approach to accurate TCR Vβ modeling by transfer learning and train a model using order of magnitude less structures (< 200 TCRs vs. > 2,000 mAbs & Nbs). The high modeling speed of NanoNet (a few milliseconds per Nb), enables accurate modeling of entire antibody repertoires from Next Generation Sequencing experiments. This will enable analysis of the serum repertoire according to antigen specificity. We applied the method for modeling a large dataset of GST Nb sequences obtained from a single animal and found that sequence similarity often corresponds with structural similarity (Fig. 6). In addition, we tested whether the NanoNet loop accuracy enables accurate antigen-Nb docking for epitope mapping.

The current NanoNet implementation is trained to predict only Cɑ coordinates. Representation of side chains (for example using center of mass) in the NanoNet, as well as refinement using GNN-based deep-learning approaches can further improve prediction accuracy ^32^. Our approach can be extended for modeling the whole mAb structure (light and heavy chains) by training a similar network for the light chain prediction and combining the light and heavy chains using most similar template structure ^33,34^. We expect that NanoNet will be applicable in therapeutic applications that require epitope mapping, as well as studies that perform serum repertoire studies. In the future, the network can be extended towards design of novel sequences with high stability and specificity for antigen of interest ^35^.

## Methods

### Network architecture

The sequences are represented by an input tensor of 140×22, where 140 represents the maximal length of the heavy chain and the 22 channels are used for one-hot encoding of the 20 amino acids (one channel for unknown amino acid and one for insertion). The tensor is padded with insertion values for all sequences shorter than 140. The network consists of two 1D ResNets ^19^ (Fig. 1A). The first ResNet has a relatively big kernel size of 25 to enable the network to learn the frame and CDR loops. Next, we convert the tensors to 140×140 dimension using a 1D convolution with 140 kernels. This second ResNet captures the inter-residue interactions and consists of kernels of size 5 with dilated convolutions ^36^. We use five different dilation values (1, 2, 4, 8, and 16). Finally, we convert the tensors to 140×70 and then to 140×3. This last tensor represents the coordinates of the Cɑ atoms and is compared directly to the actual Cɑ coordinates to calculate loss. The whole network consists of ~2,000,000 parameters. We add a dropout layer of 25% after the second ResNet to prevent overfitting. The network was implemented using the TensorFlow library with keras ^37^.

### Training

The training was performed using ADAM optimizer ^38^ with a learning rate of 0.001, batch size of 16 and ~130 epochs using a model checkpoint on the validation loss (Fig. 1C). The training took less than 10 minutes on a GeForce RTX 2080 Ti.

### Structure prediction

The prediction of Cɑ coordinates based on the trained network is straightforward and takes about 6 milliseconds on a GPU or about 20 milliseconds on a CPU. The full atom model can be obtained using PULCHRA ^39^. NanoNet produces a single structural model for each input sequence.

### Retraining for TCRs structure prediction

The NanoNet network for TCRs was trained starting from the pretrained antibody network using the TCR training set structures with the same parameters, loss function and learning rate. It was trained for 50 epochs (Fig. S1

### Dataset

Due to a relatively small number of Nb structures in the PDB, we trained our model using the heavy chains of both mAbs and Nbs. The antibody structures were obtained from abYbank/AbDb ^40^ and SAbDab ^41,42^; a total of 2,085 non-redundant structures of Nbs (319) and mAb heavy chains (1,766) were used. We selected non-redundant structures with resolution of 4Å or better. Missing residues (< 6 consecutive residues) were added by MODELLER v9.18 automodel protocol ^43,44^. The data was split into training, testing, and validation sets. In total 1,843 structures (1590 mAb, 253 Nb) were used for training and 150 structures (129 mAb, 21 Nb) were used for validation (7.5%). In addition, we used two test sets: mAb_test and Nb_test. The mAb_test consisted of 47 mAb heavy chain structures (RosettaAntibody test set ^45,46^). The Nb_test consisted of 44 Nb structures that were deposited into PDB after July 2019 with resolution higher than 2.5Å. The average CDR3 loop length was 14.0, 14.2, 12.9, and 15.3 amino acids for train, validation, mAb_test, and Nb_test, respectively (Fig. 1D). We used DeepAb data separation to enable direct comparison of mAb modeling which was divided into train and test using 99% sequence identity cutoff ^16,17^. Specifically, for assessment of Nb modeling we separated the available Nb structures such that there are no sequences with sequence identity higher than 90% between the train and test sets. The TCR structure dataset with 196 structures was obtained from the STCRDab ^47^. The data was split into train, validation, and test sets similarly to Nb dataset, this time using a 92% sequence identity cutoff which resulted in 15 structures in the test set. For TCRs, the average CDR3 loop length was 14.0 and 14.5 amino acids for train and test, respectively (Fig S1A).

### Evaluation

We evaluate the accuracy of the predicted models using the Cɑ RMSD calculated on the whole VH structure, as well as the RMSD of the frame and the three CDR loops. The RMSD of the loops was calculated based on the superposition of the whole structure that minimizes RMSD ^48^. The CDRs were defined using the IMGT numbering scheme and definition ^49^.

### Comparison to RosettaAntibody

We used the recommended RosettaAntibody protocol for mAb heavy chains and for Nbs ^8,50^. Identical structures were excluded from RosettaAntibody template search. In total, 100 loops were generated and the best-scoring was selected for comparison.

### Docking and modeling of SARS-CoV-2 RBD nanobodies

The docking was done using PatchDock antibody-antigen protocol ^51,52^ that focuses the search on the CDR loops.

### Sequence embedding

The sequences were first aligned using ANARCI antibody numbering tool ^53^ that numbers the sequence based on 126 canonical positions. After the alignment, the sequences were converted into a one-hot encoding vector representation, resulting in a representation with a vector of length 2,646 for each sequence.

### Structural distance between the Nbs

Structural similarity between a pair of Nb structures was defined by counting the number of pairs of Cɑ atoms (one from each Nb structure) within a short distance (<1Å). Note that NanoNet predicts aligned Nb structures, therefore the number of aligned Cɑ atoms is a measure of structural similarity. We define a structural distance as follows:

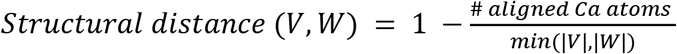

### Availability

The source code, the trained model and a Colaboratory notebook for running NanoNet are available at GitHub: https://github.com/dina-lab3D/NanoNet

