## Supplementary material for "NanoNet: Rapid end-to-end nanobody modeling by deep learning at sub angstrom resolution"

**Table S1.** NanoNet results for the Nb test set. RMSDs for the whole VHH domain, followed by frame and CDRs1-3, CDR3 length, and maximal sequence identity to an Nb in the training set.

| PDB | VHH | Fr | CDR1 | CDR2 | CDR3 | CDR3 Length | Max train identity |
|---|---|---|---|---|---|---|---|
| **6ocd** | 2.58 | 2.76 | 1.31 | 1.63 | 2.48 | 16 | 0.84 |
| **6z6v** | 1.13 | 0.55 | 1.68 | 1.08 | 2.28 | 19 | 0.79 |
| **7a4t** | 1.31 | 1.01 | 2.73 | 0.91 | 1.81 | 14 | 0.89 |
| **7c8v** | 0.82 | 0.58 | 1.49 | 1.72 | 0.80 | 8 | 0.89 |
| **6yu8** | 1.89 | 0.90 | 3.25 | 2.13 | 4.01 | 15 | 0.83 |
| **6xw5** | 1.42 | 0.97 | 3.37 | 1.14 | 2.15 | 11 | 0.84 |
| **7n0i** | 1.38 | 1.14 | 1.43 | 1.24 | 2.20 | 19 | 0.85 |
| **6waq** | 1.19 | 0.72 | 2.17 | 0.62 | 2.23 | 18 | 0.89 |
| **6xyf** | 1.55 | 0.83 | 3.42 | 0.85 | 2.70 | 19 | 0.85 |
| **7a0v** | 1.18 | 0.63 | 2.78 | 0.46 | 2.10 | 16 | 0.82 |
| **7d30** | 2.41 | 1.14 | 2.57 | 2.25 | 5.90 | 14 | 0.88 |
| **7mfv** | 0.85 | 0.46 | 2.05 | 1.52 | 1.14 | 8 | 0.88 |
| **7a50** | 1.65 | 0.80 | 1.61 | 2.53 | 3.44 | 19 | 0.86 |
| **7ldj** | 2.03 | 1.11 | 3.31 | 1.59 | 3.69 | 23 | 0.88 |
| **7kn5** | 2.17 | 0.91 | 3.64 | 0.88 | 4.38 | 22 | 0.87 |
| **6uht** | 3.38 | 3.43 | 5.14 | 1.09 | 2.46 | 18 | 0.83 |
| **7kjh** | 1.39 | 1.04 | 2.05 | 2.09 | 2.21 | 13 | 0.87 |
| **6xxo** | 1.60 | 1.18 | 3.62 | 2.70 | 1.29 | 17 | 0.78 |
| **7aqg** | 1.48 | 0.76 | 1.43 | 3.85 | 2.33 | 14 | 0.86 |
| **6x05** | 2.34 | 1.31 | 3.96 | 2.46 | 4.62 | 15 | 0.75 |
| **6xw7** | 0.73 | 0.45 | 0.70 | 0.83 | 1.85 | 10 | 0.86 |
| **6lz2** | 1.29 | 0.60 | 2.26 | 1.47 | 3.53 | 8 | 0.88 |
| **6z10** | 1.63 | 0.97 | 1.79 | 0.49 | 3.22 | 22 | 0.81 |
| **6obm** | 2.08 | 1.30 | 4.43 | 1.25 | 3.73 | 14 | 0.88 |
| **6lr7** | 1.07 | 0.50 | 2.23 | 0.57 | 1.99 | 20 | 0.78 |
| **6xzu** | 2.50 | 1.18 | 2.57 | 1.72 | 6.74 | 12 | 0.88 |
| **7mfu** | 1.38 | 0.66 | 1.34 | 4.50 | 0.58 | 8 | 0.88 |
| **6xw4** | 2.06 | 1.23 | 4.66 | 2.20 | 3.33 | 15 | 0.83 |
| **6obg** | 1.79 | 0.96 | 2.76 | 1.09 | 4.00 | 14 | 0.86 |
| **6zrv** | 1.67 | 0.77 | 1.59 | 1.99 | 3.85 | 16 | 0.90 |
| **7now** | 1.50 | 0.76 | 2.84 | 0.68 | 2.83 | 21 | 0.89 |
| **6obo** | 1.28 | 0.66 | 2.66 | 0.69 | 2.61 | 14 | 0.89 |
| **7k84** | 1.82 | 0.78 | 1.05 | 1.68 | 4.12 | 19 | 0.84 |
| **6oca** | 2.18 | 1.15 | 3.10 | 4.59 | 3.04 | 18 | 0.82 |
| **6ui1** | 2.19 | 1.19 | 2.11 | 1.39 | 4.79 | 18 | 0.76 |
| **7a48** | 1.17 | 0.69 | 2.16 | 1.37 | 2.44 | 11 | 0.84 |
| **6obc** | 1.41 | 0.88 | 2.57 | 1.09 | 2.83 | 13 | 0.76 |
| **6z1v** | 1.34 | 0.73 | 1.41 | 0.86 | 3.08 | 16 | 0.83 |
| **7kgj** | 1.84 | 0.79 | 1.92 | 4.11 | 3.63 | 14 | 0.89 |
| **6xw6** | 1.22 | 0.69 | 2.45 | 1.67 | 1.88 | 21 | 0.82 |
| **6obe** | 1.92 | 1.21 | 3.79 | 2.21 | 3.45 | 13 | 0.79 |
| **7d2z** | 1.50 | 1.50 | 0.67 | 2.24 | 0.99 | 8 | 0.89 |
| **7n0r** | 2.53 | 1.09 | 1.49 | 2.36 | 6.57 | 14 | 0.87 |
| **7lvu** | 1.38 | 0.81 | 2.87 | 2.66 | 1.78 | 16 | 0.86 |
| **Mean** | **1.66** | **1.00** | **2.46** | **1.74** | **2.98** | **15.30** | **0.85** |
| **STD** | **0.54** | **0.53** | **1.04** | **1.03** | **1.38** | **4.07** | **0.04** |
| **Median** | **1.52** | **0.89** | **2.35** | **1.56** | **2.76** | **15.00** | **0.86** |

**Table S2.** NanoNet results on the mAb test set. RMSDs for the whole VH domain, followed by frame and CDRs1-3, CDR3 length, and maximal sequence identity to a mAb VH in the training set.

| PDB | VH | FR | CDR1 | CDR2 | CDR3 | CDR3 Length | Max train identity |
|---|---|---|---|---|---|---|---|
| **1mtq** | 0.49 | 0.34 | 0.33 | 0.40 | 1.19 | 11 | 0.90 |
| **2r8s** | 1.29 | 0.61 | 0.83 | 2.73 | 2.69 | 14 | 0.96 |
| **3nps** | 1.38 | 0.87 | 1.55 | 1.03 | 2.73 | 19 | 0.95 |
| **2v17** | 1.00 | 0.74 | 0.40 | 0.78 | 2.23 | 13 | 0.88 |
| **1seq** | 1.42 | 0.53 | 1.03 | 0.87 | 3.59 | 16 | 0.88 |
| **1fns** | 1.01 | 0.63 | 0.50 | 0.51 | 2.32 | 16 | 0.91 |
| **3go1** | 1.25 | 0.50 | 1.67 | 1.98 | 2.70 | 16 | 0.75 |
| **2adf** | 0.59 | 0.34 | 0.41 | 0.60 | 1.56 | 11 | 0.91 |
| **1mfa** | 0.65 | 0.55 | 0.47 | 0.47 | 1.30 | 11 | 0.97 |
| **3eo9** | 0.92 | 0.58 | 0.49 | 0.89 | 2.13 | 14 | 0.87 |
| **4nzu** | 1.77 | 0.64 | 1.39 | 0.79 | 4.30 | 18 | 0.80 |
| **2e27** | 1.06 | 0.48 | 0.88 | 0.37 | 3.37 | 9 | 0.89 |
| **3gnm** | 0.64 | 0.49 | 0.44 | 0.38 | 1.45 | 11 | 0.85 |
| **2ypv** | 1.16 | 1.01 | 0.35 | 0.76 | 2.27 | 12 | 0.84 |
| **3p0y** | 0.91 | 0.54 | 1.02 | 0.85 | 2.06 | 14 | 0.90 |
| **3t65** | 0.46 | 0.33 | 0.45 | 0.60 | 0.91 | 13 | 0.95 |
| **2xwt** | 0.80 | 0.50 | 1.20 | 0.51 | 1.84 | 12 | 0.87 |
| **3umt** | 0.76 | 0.45 | 0.51 | 0.60 | 1.93 | 12 | 0.88 |
| **1dlf** | 1.37 | 0.58 | 1.38 | 0.97 | 3.76 | 12 | 0.91 |
| **3e8u** | 0.81 | 0.57 | 0.84 | 0.46 | 1.98 | 10 | 0.83 |
| **3mxw** | 0.92 | 0.75 | 1.08 | 1.05 | 1.59 | 12 | 0.86 |
| **3mlr** | 1.90 | 1.04 | 1.02 | 0.66 | 4.43 | 17 | 0.75 |
| **4h20** | 0.97 | 0.59 | 2.04 | 0.61 | 1.90 | 12 | 0.87 |
| **1gig** | 1.06 | 0.58 | 0.90 | 0.52 | 2.47 | 16 | 0.88 |
| **3liz** | 1.31 | 0.41 | 0.58 | 0.28 | 3.92 | 12 | 0.87 |
| **2vxv** | 1.24 | 0.49 | 0.54 | 1.00 | 3.32 | 14 | 0.89 |
| **3g5y** | 0.42 | 0.34 | 0.67 | 0.35 | 0.72 | 9 | 0.96 |
| **3lmj** | 2.44 | 0.75 | 3.51 | 6.56 | 3.71 | 18 | 0.83 |
| **4h0h** | 0.91 | 0.82 | 0.34 | 0.37 | 1.72 | 12 | 0.88 |
| **3oz9** | 0.88 | 0.43 | 0.35 | 0.81 | 2.39 | 12 | 0.89 |
| **3giz** | 0.77 | 0.39 | 0.47 | 0.37 | 1.92 | 15 | 0.91 |
| **1oaq** | 0.69 | 0.52 | 0.51 | 0.55 | 1.46 | 13 | 0.95 |
| **2d7t** | 0.93 | 0.79 | 0.66 | 0.37 | 2.04 | 9 | 0.93 |
| **1jfq** | 0.74 | 0.78 | 0.34 | 0.65 | 0.63 | 14 | 0.98 |
| **3hc4** | 0.93 | 0.93 | 0.40 | 0.39 | 1.45 | 9 | 0.86 |
| **1mlb** | 0.48 | 0.35 | 0.20 | 0.48 | 1.23 | 9 | 0.96 |
| **2w60** | 0.65 | 0.46 | 0.53 | 0.31 | 1.58 | 11 | 0.92 |
| **2fb4** | 1.37 | 0.57 | 0.56 | 0.70 | 3.24 | 19 | 0.82 |
| **2fbj** | 0.70 | 0.58 | 0.51 | 0.36 | 1.49 | 11 | 0.90 |
| **3vow** | 0.65 | 0.57 | 0.78 | 0.77 | 0.94 | 13 | 0.84 |
| **1jpt** | 0.75 | 0.59 | 0.31 | 0.42 | 1.79 | 10 | 0.83 |
| **4f57** | 1.16 | 0.43 | 0.68 | 1.45 | 2.70 | 18 | 0.79 |
| **3i9g** | 1.35 | 0.46 | 0.55 | 0.52 | 3.74 | 14 | 0.80 |
| **1nlb** | 0.58 | 0.57 | 0.43 | 0.52 | 0.81 | 11 | 0.91 |
| **3hnt** | 0.49 | 0.36 | 0.30 | 0.31 | 1.18 | 11 | 0.88 |
| **3m8o** | 1.66 | 0.87 | 1.19 | 1.36 | 4.78 | 10 | 0.76 |
| **4hpy** | 1.13 | 0.38 | 2.98 | 0.70 | 2.22 | 13 | 0.93 |
| **Mean** | **0.99** | **0.58** | **0.82** | **0.83** | **2.25** | **12.94** | **0.88** |
| **STD** | **0.41** | **0.18** | **0.66** | **0.96** | **1.05** | **2.79** | **0.06** |
| **Median** | **0.92** | **0.57** | **0.55** | **0.60** | **2.04** | **12.00** | **0.88** |

**Table S3.** NanoNet results for the TCR test set. RMSDs for the whole Vβ domain, followed by frame and CDRs1-3, CDR3 length, and maximal sequence identity to an Nb in the training set.

| PDB | Vβ | Fr | CDR1 | CDR2 | CDR3 | CDR3 Length | Max train identity |
|---|---|---|---|---|---|---|---|
| **6vrn** | 0.99 | 0.76 | 0.47 | 0.56 | 2.02 | 15 | 0.92 |
| **6c61** | 1.76 | 1.73 | 0.60 | 0.80 | 2.65 | 16 | 0.90 |
| **2ial** | 0.72 | 0.77 | 0.44 | 0.57 | 0.75 | 11 | 0.93 |
| **6fr8** | 1.05 | 0.61 | 0.55 | 0.45 | 2.51 | 14 | 0.93 |
| **4ww1** | 1.02 | 0.90 | 1.12 | 0.71 | 1.71 | 12 | 0.82 |
| **1kgc** | 0.87 | 0.67 | 0.62 | 0.67 | 1.79 | 13 | 0.81 |
| **6ovn** | 1.80 | 1.66 | 0.85 | 2.31 | 2.40 | 17 | 0.93 |
| **4ei6** | 2.46 | 2.69 | 1.82 | 1.87 | 1.80 | 14 | 0.71 |
| **6r2l** | 2.12 | 0.95 | 0.83 | 5.01 | 3.12 | 18 | 0.86 |
| **2cdf** | 1.23 | 0.98 | 0.78 | 0.64 | 2.39 | 15 | 0.93 |
| **6frb** | 0.82 | 0.76 | 0.65 | 0.96 | 1.13 | 12 | 0.64 |
| **6lir** | 1.04 | 1.12 | 0.60 | 1.07 | 0.81 | 14 | 0.58 |
| **6ovo** | 1.81 | 0.93 | 1.16 | 1.00 | 4.07 | 17 | 0.93 |
| **3vxq** | 0.85 | 0.74 | 0.52 | 0.44 | 1.59 | 13 | 0.93 |
| **5d2n** | 1.07 | 0.82 | 0.65 | 0.87 | 2.07 | 16 | 0.92 |
| **Mean** | **1.31** | **1.07** | **0.78** | **1.20** | **2.05** | **14.47** | **0.85** |
| **STD** | **0.54** | **0.55** | **0.36** | **1.18** | **0.87** | **2.07** | **0.12** |
| **Median** | **1.05** | **0.90** | **0.65** | **0.80** | **2.02** | **14.00** | **0.92** |

**Table S4.** Docking results for NanoNet generated models. Minimal ligand and interface RMSDs for docking models generated by PatchDock.

| PDB | Antigen | Min ligand | Min interface |
|---|---|---|---|
| **6yu8** | Ebola RNA methyltransferase | 6.52 | 4.25 |
| **6xw5** | MNV capsid protein P-domain | 5.04 | 2.54 |
| **6xw7** | MNV capsid protein P-domain | 4.07 | 1.76 |
| **6xw4** | MNV capsid protein P-domain | 9.98 | 3.17 |
| **6xw6** | MNV capsid protein P-domain | 4.72 | 2.13 |
| **6waq** | SARS-CoV-1 RBD | 4.31 | 2.00 |
| **7n0i** | SARS-CoV-2 N protein | 5.22 | 1.67 |
| **7n0r** | SARS-CoV-2 N protein | 6.22 | 4.17 |
| **7c8v** | SARS-CoV-2 RBD | 2.39 | 3.76 |
| **7d30** | SARS-CoV-2 RBD | 4.95 | 2.34 |
| **7ldj** | SARS-CoV-2 RBD | 8.72 | 5.46 |
| **7kn5** | SARS-CoV-2 RBD | 5.01 | 3.92 |
| **7mfu** | SARS-CoV-2 RBD | 4.02 | 3.52 |
| **7kgj** | SARS-CoV-2 RBD | 5.58 | 5.25 |
| **7d2z** | SARS-CoV-2 RBD | 6.27 | 4.63 |
| **nb105** | SARS-CoV-2 RBD | 6.65 | 2.62 |
| **nb21** | SARS-CoV-2 RBD | 3.81 | 5.07 |
| **Mean** | - | **5.50** | **3.43** |
| **STD** | - | **1.83** | **1.26** |
| **Median** | - | **5.04** | **3.52** |

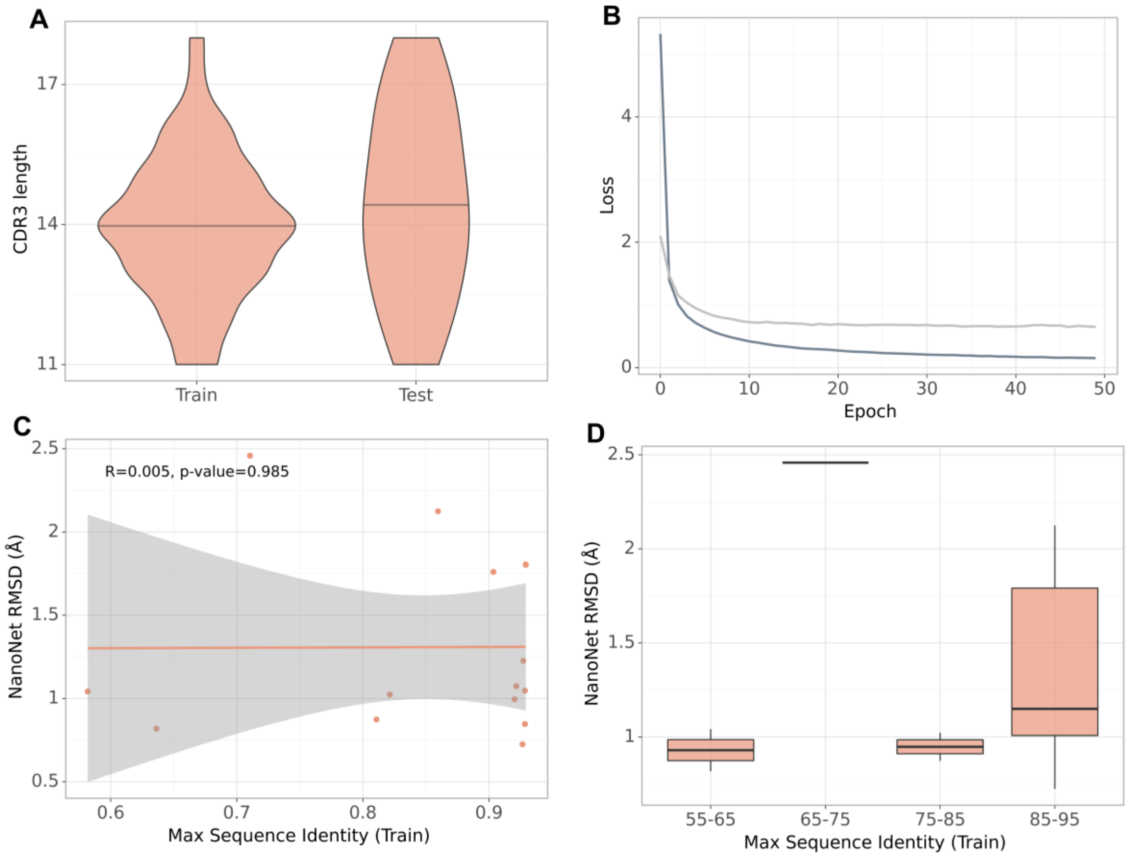


**Fig S1.** TCR β chain test set. A. CDR3 lengths of the training and test sets. B. Training and validation loss during the training process. C. Test set performance (measured as Vβ RMSD) as a function of maximal sequence identity to the train set, each dot represents a structure from the test set. D. Test set performance with boxplots for sequence identity ranges.

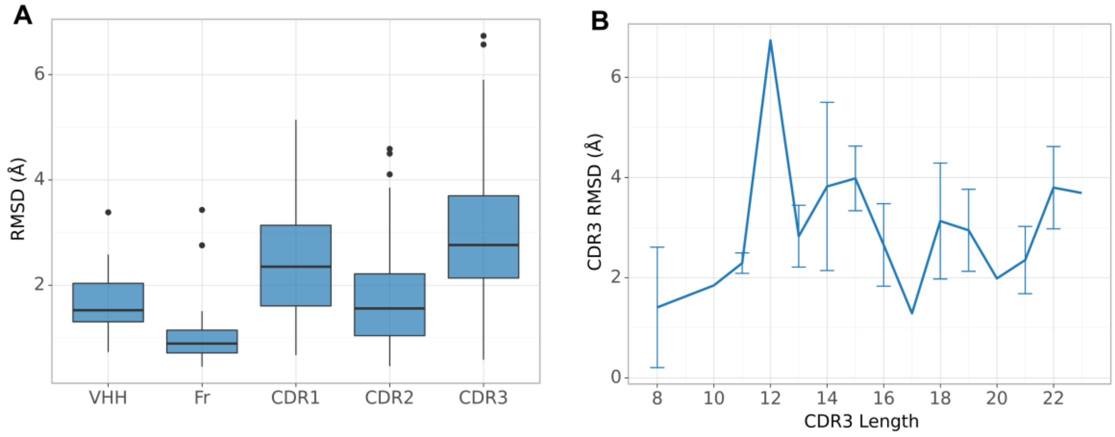


**Fig S2**: NanoNet results for the entire Nb test set (44 structures). A. Boxplots of RMSDs of the whole VHH region, framework, CDR1-3 loops. B. RMSD of CDR3 loop as a function of loop length.

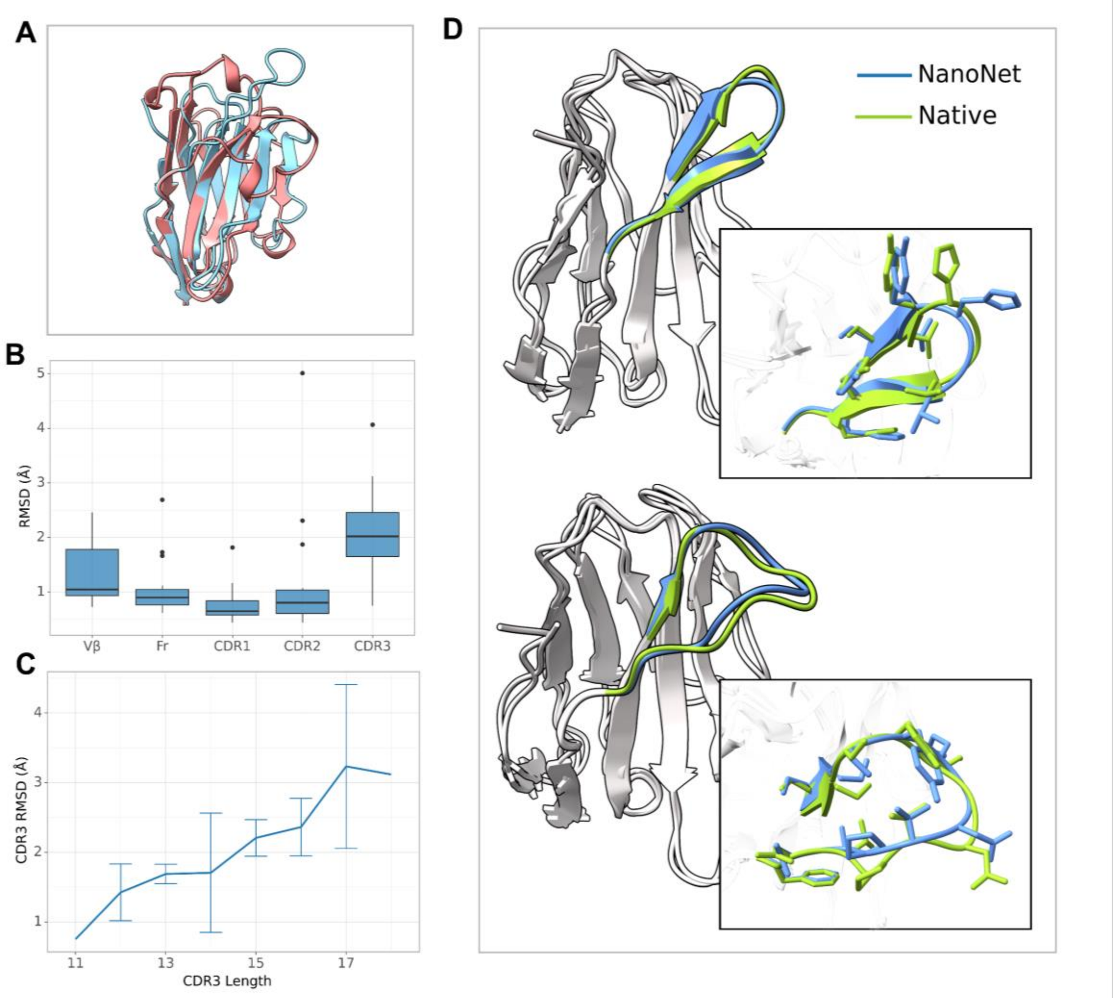


**Fig. S3**: TCR β chain modeling. A. TCR Vβ (PDB 6r2l, blue) aligned to an Nb (PDB 6uht, red) B. Boxplots of RMSDs of the whole Vβ region, framework, CDR1-3 loops for the test set TCRs. C. RMSD of CDR3 loop as a function of loop length on the test set of 15 TCRs. D. Test set examples of modeled structure (blue) vs. experimental (green): PDB 2ial (top) and 6lir (bottom), RMSDs 0.72Å and 1.0Å, respectively.
